## Supplemental Table 1 for "CM1-driven assembly and activation of Yeast γ-Tubulin Small Complex underlies microtubule nucleation"

| | Xrcc4-Spc110p$^{164-207}$ |
|---|---|
| PDB ID | 7M3P |
| Beam line | Advanced Light Source beamline 8.3.1 |
| Wavelength | 1.116 |
| Resolution range | 35.73-2.0 (2.071-2.0) |
| Space group | P 1 |
| Unit cell a b γ | 38.131 52.168 55.916 |
| Unit cell α β γ | 100.804 94.148 112.831 |
| Total reflections | 50099 (5015) |
| Unique reflections | 24292 (2436) |
| Multiplicity | 2.1 (2.1) |
| Completeness (%) | 93 (94) |
| Mean I/sigma(I) | 17.36 (5.45) |
| Wilson B-factor | 27.9 |
| R-merge | 0.03095 (0.1635) |
| R-meas | 0.0426 (0.2245) |
| CC1/2 | 0.999 (0.939) |
| CC* | 1 (0.984) |
| Reflections used in refinement | 24291 (2436) |
| Reflections used for R-free | 1462 (152) |
| R-work | 0.2043 (0.2643) |
| R-free | 0.2470 (0.3016) |
| CC (work) | 0.947 (0.847) |
| CC (free) | 0.926 (0.710) |
| Number of non-hydrogen atoms | 2965 |
| macromolecules | 2711 |
| solvent | 254 |
| Protein residues | 332 |
| RMS (bonds) | 0.005 |
| RMS (angles) | 0.58 |
| Ramachandran favored (%) | 97 |
| Ramachandran allowed (%) | 3.1 |
| Ramachandran outliers (%) | 0 |
| Rotamer outliers (%) | 0 |
| Clashscore | 3.34 |
| Average B-factor | 44.25 |
| macromolecules | 44.38 |
| solvent | 42.94 |
| Number of TLS groups | 24 |

**Table S1: X-ray data collection and refinement statistics.**
