## Supplemental Table 2 for "CM1-driven assembly and activation of Yeast γ-Tubulin Small Complex underlies microtubule nucleation"

| Name | γTuSC monomer | γTuSC dimer | γTuRC$^{ss}$ | γTuRC$^{WT}$ open | γTuRC$^{WT}$ closed |
|---|---|---|---|---|---|
| PDB ID | 7M2Z | | 7M2W | 7M2X | 7M2Y |
| EMDB ID | EMD-23638 | EMD-23639 | EMD-23635 | EMD-23636 | EMD-23637 |
| | | | **Data collection/processing** | | |
| Microscope | Polara | Polara | Krios | Krios | Krios |
| Camera | K2 | K2 | K2 | K2 | K2 |
| Magnification | 39891 | 39891 | 47214 | 47214 | 47214 |
| Voltage (kV) | 300 | 300 | 300 | 300 | 300 |
| Total electron dose ($e^-/Å^2$) | 81 | 81 | 72 | 80 | 80 |
| Defocus range (µm) | 0.4-5.2 | 0.4-5.2 | 0.5-2.5 | 0.6-3.0 | 0.6-3.0 |
| Processing pixel size (Å) | 0.6267 | 0.6267 | 1.059 | 1.4183 | 1.4183 |
| Pixel size (Å) Physical/(Super-resolution) | 1.2534/ (0.6267) | 1.2534/ (0.6267) | 1.059/ (0.5295) | 1.059/ (0.5295) | 1.059/ (0.5295) |
| Micrographs | 7381 | 7381 | 3024 | 2204 | 2204 |
| Final particles | 398841 | 506014 | 148911 | 36251 | 20420 |
| Symmetry | C1 | C1 | C1 | C1 | C1 |
| High resolution limit used in refinement | 6.0 | 8.0 | 3.7 | 4.5 | 4.5 |
| Resolution Å (FSC 0.143) | 3.7 | 4.5 | 3.0 | 3.6 | 4.1 |
| | | | **Model Composition** | | |
| Non-hydrogen atoms | 18332 | | 39371 | 18706 | 18563 |
| Protein residues | 2253 | | 4827 | 2397 | 2278 |
| Ligands | 2 | | 4 | 2 | 2 |
| | | | **Model Refinement** | | |
| Map-model FSC (0.5) | 4.2 | | 3.4 | 4.2 | 4.5 |
| RMSD bond lengths (Å) | 0.007 | | 0.005 | 0.022 | 0.022 |
| RMSD bond angles (degrees) | 1.141 | | 1.011 | 1.760 | 1.793 |
| | | | **Validation** | | |
| MolProbity score | 2.21 | | 1.91 | 1.09 | 1.25 |
| Clash score | 17.15 | | 12.2 | 2.29 | 3.66 |
| C-beta deviations (%) | 0 | | 0.02 | 0 | 0 |
| Rotamer outliers (%) | 0 | | 0.23 | 0.14 | 0.1 |
| Ramachandran favored (%) | 92.24 | | 95.5 | 97.62 | 97.56 |
| Ramachandran disallowed (%) | 0.04 | | 0.02 | 0.35 | 0.27 |

**Table S2. CryoEM data collection, processing, and modeling statistics.**
