## Supplemental Figures for "CM1-driven assembly and activation of Yeast γ-Tubulin Small Complex underlies microtubule nucleation"

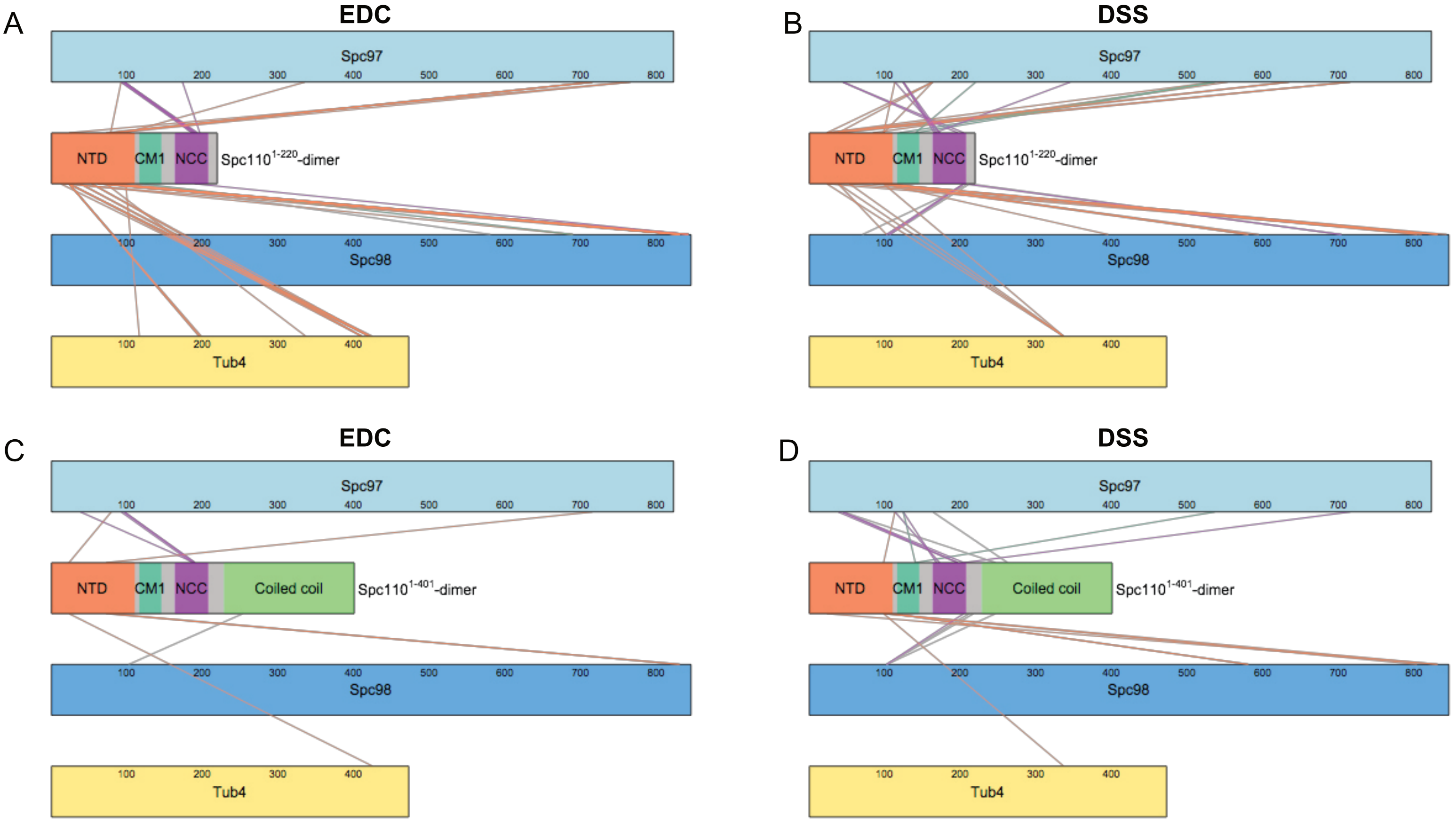


E

| Crosslinker | Position | | | Spc110 construct | |
|---|---|---|---|---|---|
| | Spc110 | Spc97 | Spc98 | GCN4 dimer | 1-401-GST |
| EDC | 187* | 93 | | + | + |
| | 187 | 97 | | + | |
| | 190* | 93 | | + | |
| | 190* | 97 | | + | + |
| | 190 | | 843 | + | |
| | 191 | 97 | | + | + |
| | 192* | 97 | | + | |
| | 194 | 97 | | + | |
| | 197 | 174 | | + | |
| DSS | 166* | 125 | | + | |
| | 166 | 346 | | + | |
| | 169* | 125 | | + | + |
| | 173* | 114 | | + | + |
| | 173* | 125 | | + | |

**Figure 2 - figure supplement 1: Overview of XL-MS datasets.**

XL-MS datasets for γTuSC crosslinked to Spc110p$^{1-220}$-GCN4 dimer (A-B) or Spc110p$^{1-401}$-GST (C-D) with either EDC (A, C) or DSS (B, D). Each colored rectangles represents a protein in the crosslinked sample. The lengths of the rectangles are proportional to the number of residues in each protein. Crosslinks between γTuSC components and Spc110p$^{1-111}$ , Spc110p$^{CM1(117-146)}$ , and Spc110p$^{NCC\ (164-208)}$ regions are color coded orange, green, and purple, respectively.
E. Table of crosslinks between Spc110p$^{NCC(164-208)}$ and γTuSC. Spc110p crosslinks marked with an asterisk (*) are satisfied on either monomer within Spc110p$^{NCC(164-208)}$.

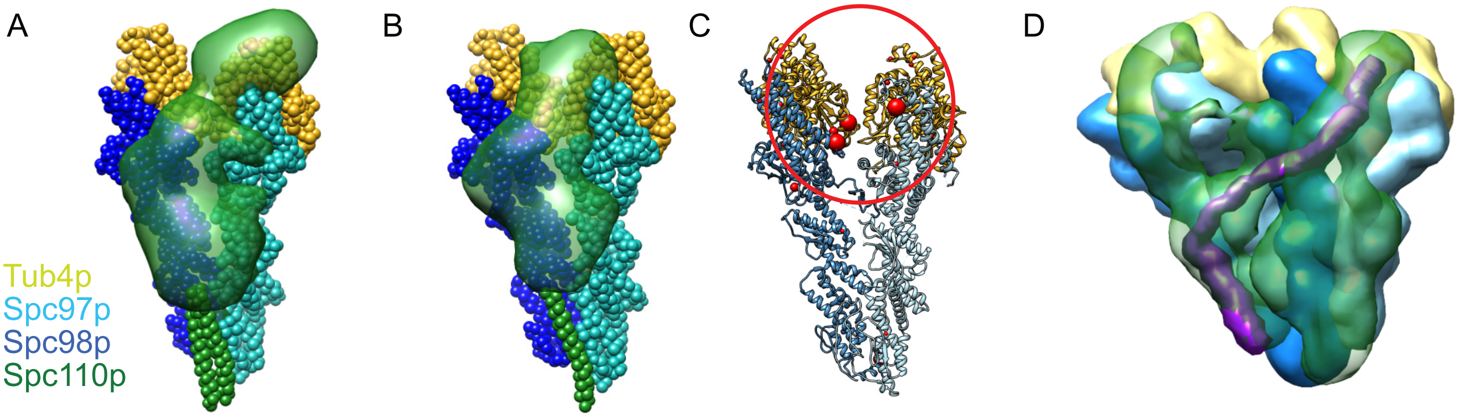


**Figure 2 - figure supplement 2: Results of integrative modeling of the Spc110p-γTuSC complex.**

*The modeling results shown are based on the γTuSC-Spc110p$^{1\text{-}220}$ GCN4 crosslinks; similar results were obtained in all cases using γTuSC-Spc110p$^{1\text{-}401}$-GST crosslinks (see Supplementary Methods)*.
A. Monomer of Spc110p$^{1\text{-}220}$-GCN4 bound to γTuSC. γTuSC and Spc110p NCC, which were kept fixed in the modeling, are shown as spherical beads with one bead per residue. Spc110p$^{1\text{-}163}$ is shown as a localization density map, showing the positions of the Spc110p NTD in the ensemble of models from the top cluster.
B. Dimer of Spc110p$^{1\text{-}220}$-GCN4 bound to γTuSC, with a representation similar to (A).
C. A γTuSC monomer with EDC and DSS crosslinks from the NTD (residues 3-113) of the Spc110p$^{1\text{-}220}$-GCN4 dimer construct mapped onto it (Note that residues 3-113 in this construct correspond to residues 1-111 in Spc110). Each red sphere represents a residue in TuSC where the Spc110p$^{1\text{-}111}$ crosslinks. The size of the red sphere indicates the number of residues in Spc110p$^{1\text{-}111}$ that crosslink there. The smallest sphere indicates 1 residue from Spc110p$^{1\text{-}111}$ crosslinks to that γTuSC residue; the largest spheres show where 4 or more residues from Spc110p$^{1\text{-}111}$ crosslink. The red circle highlights a cluster of crosslink sites between Spc110p$^{1\text{-}111}$ and Spc97p or Spc98p at the intra-γTuSC interface.
D. Dimer of Spc110p$^{1\text{-}220}$-GCN4 bound to adjacent γTuSCs, shown as a localization density map, similar to Fig. 1D. Maps for all components are contoured at 2.5% of their respective maximum voxel values. We highlight in purple a hypothetical path integrating our modeling and structural work showing how Spc110p bridges the inter-γTuSC interface and reaches to the adjacent γTuSC where it crosslinks to the N-terminal regions of Spc97p and Spc98p at the intra-γTuSC interface.

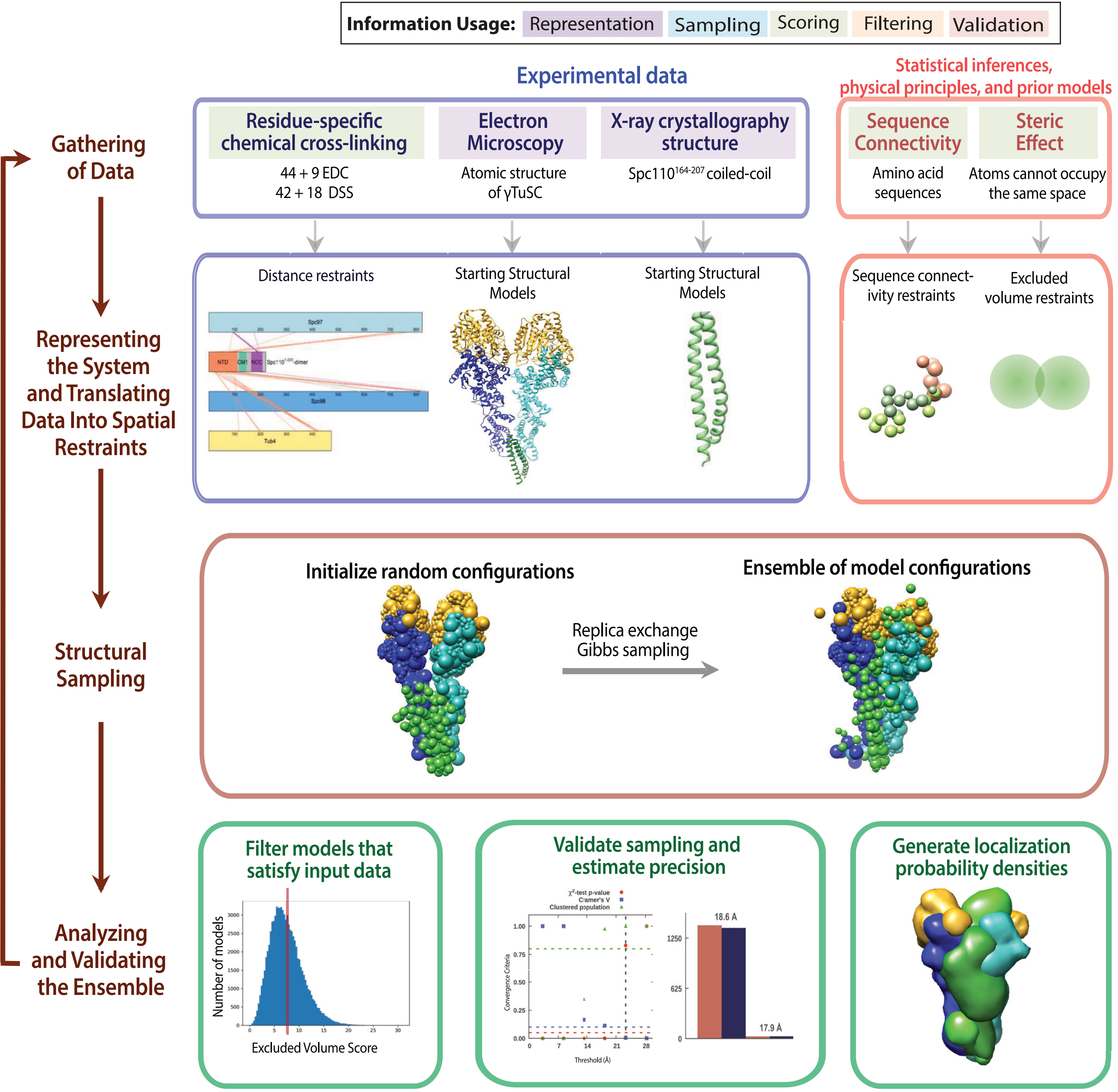


**Figure 2 - figure supplement 3: The four stages of integrative modeling of the Spc110-γTuSC complex.**

This schematic describes the integrative structure modeling procedures used in this paper. The first row details the information to be used in modeling. The background color of each information source indicates where the information is applied in modeling, as detailed in the key at the top. The second row describes how each information source is converted into spatial restraints. The third row details the sampling protocol. The last row details the analysis and validation steps of the modeling.

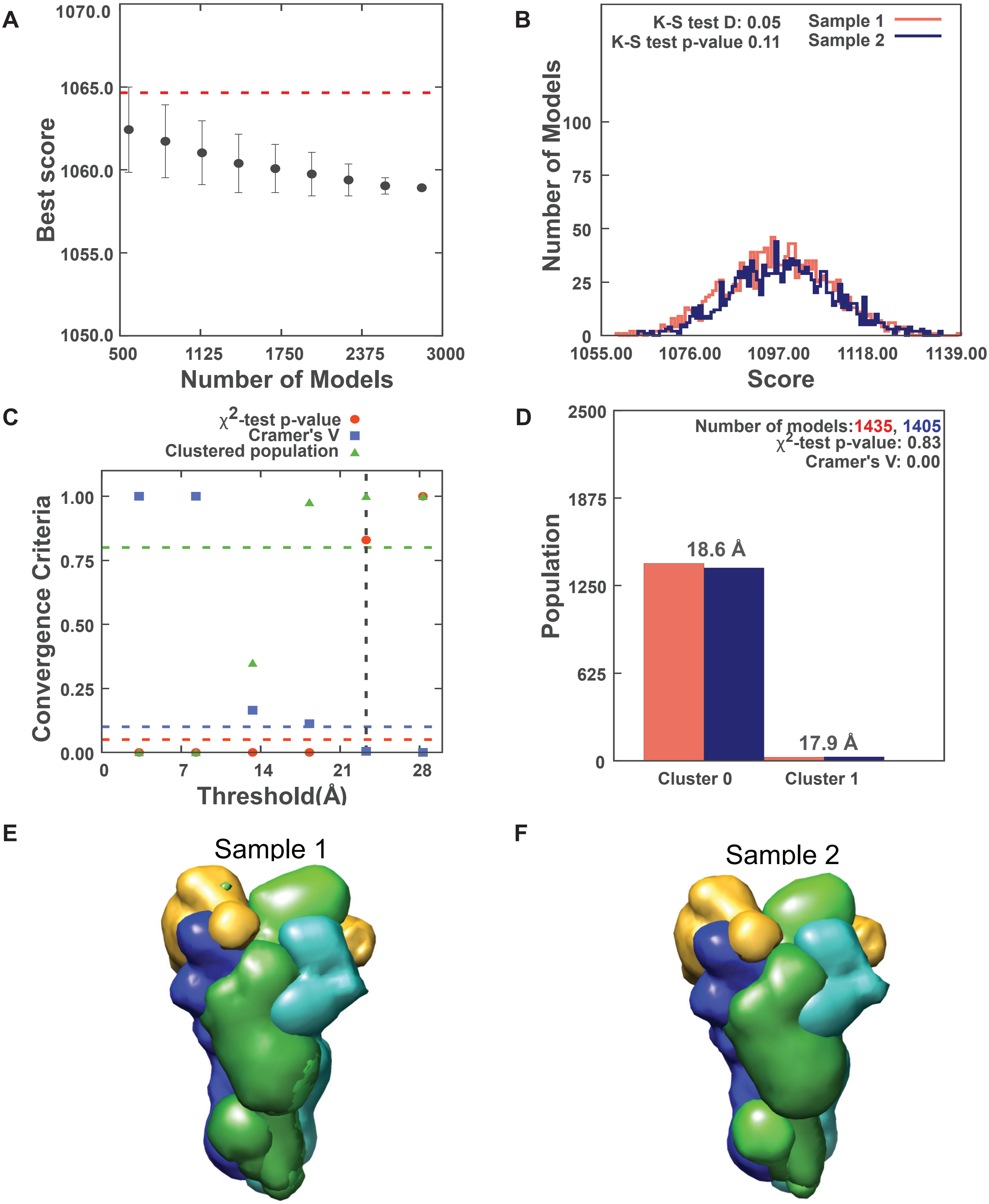


**Figure 2 - figure supplement 4: Results for sampling exhaustiveness protocol for modeling the complex of Spc110p$^{1\text{-}220}$-GCN4 dimer with γTuSC.**

A. Results of test 1, convergence of the model score, for the 2840 good-scoring models; the scores do not continue to improve as more models are computed essentially independently. The error bar represents the standard deviations of the best scores, estimated by repeating sampling of models 10 times. The red dotted line indicates a lower bound reference on the total score. B. Results of test 2, testing similarity of model score distributions between samples 1 (red) and 2 (blue); the difference in distribution of scores is not significant (Kolmogorov-Smirnov two-sample test p-value greater than 0.05) and the magnitude of the difference is small (the Kolmogorov-Smirnov two-sample test statistic D is 0.05); thus, the two score distributions are effectively equal. C. Results of test 3, three criteria for determining the sampling precision (Y-axis), evaluated as a function of the RMSD clustering threshold (X-axis). First, the p-value is computed using the χ2 -test for homogeneity of proportions (red dots). Second, an effect size for the χ2 -test is quantified by the Cramer's V value (blue squares). Third, the population of models in sufficiently large clusters (containing at least 10 models from each sample) is shown as green triangles. The vertical dotted grey line indicates the RMSD clustering threshold at which three conditions are satisfied (p-value > 0.05 [dotted red line], Cramer's V < 0.10 [dotted blue line], and the population of clustered models > 0.80 [dotted green line]), thus defining the sampling precision of 23.3 Å. D. Populations of sample 1 and 2 models in the clusters obtained by threshold-based clustering using the RMSD threshold of 23.3 Å. Cluster precision is shown for each cluster. E. and F. Results of test 4: comparison of localization probability densities of models from sample A and sample B for the major cluster (98.1% population). The cross-correlation of the density maps of the two samples is 0.99 for the Spc110 maps (green).

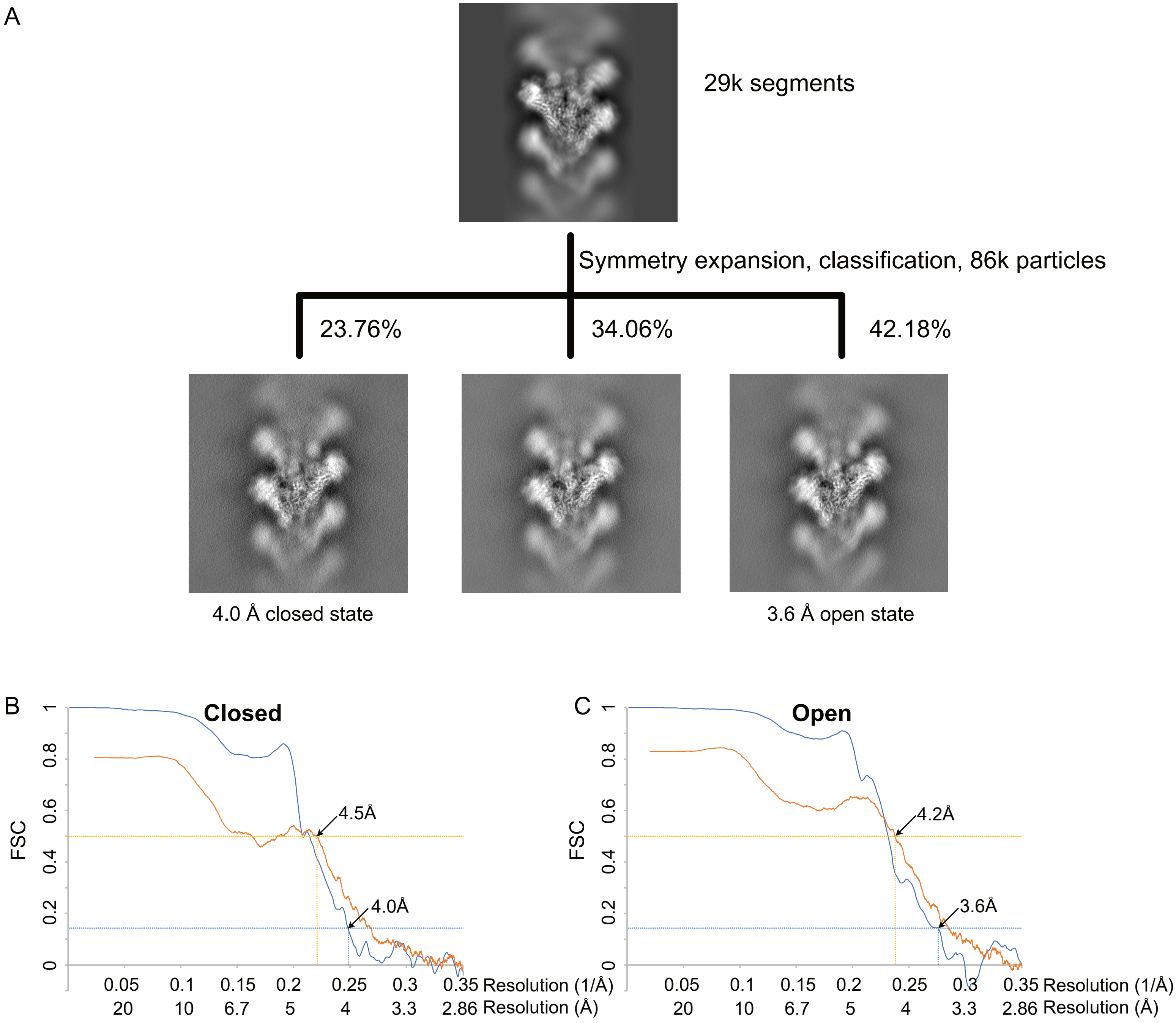


**Figure 3 - figure supplement 1: γTuRC$^{WT}$ processing and resolution**

A. Classification scheme for γTuRC$^{WT}$ processing. Images are projection images of the 3D classes obtained.
B. FSC (blue) and map to model FSC (orange) for the closed γTuRC$^{WT}$.
C. FSC (blue) and map to model FSC (orange) for the open γTuRC$^{WT}$.

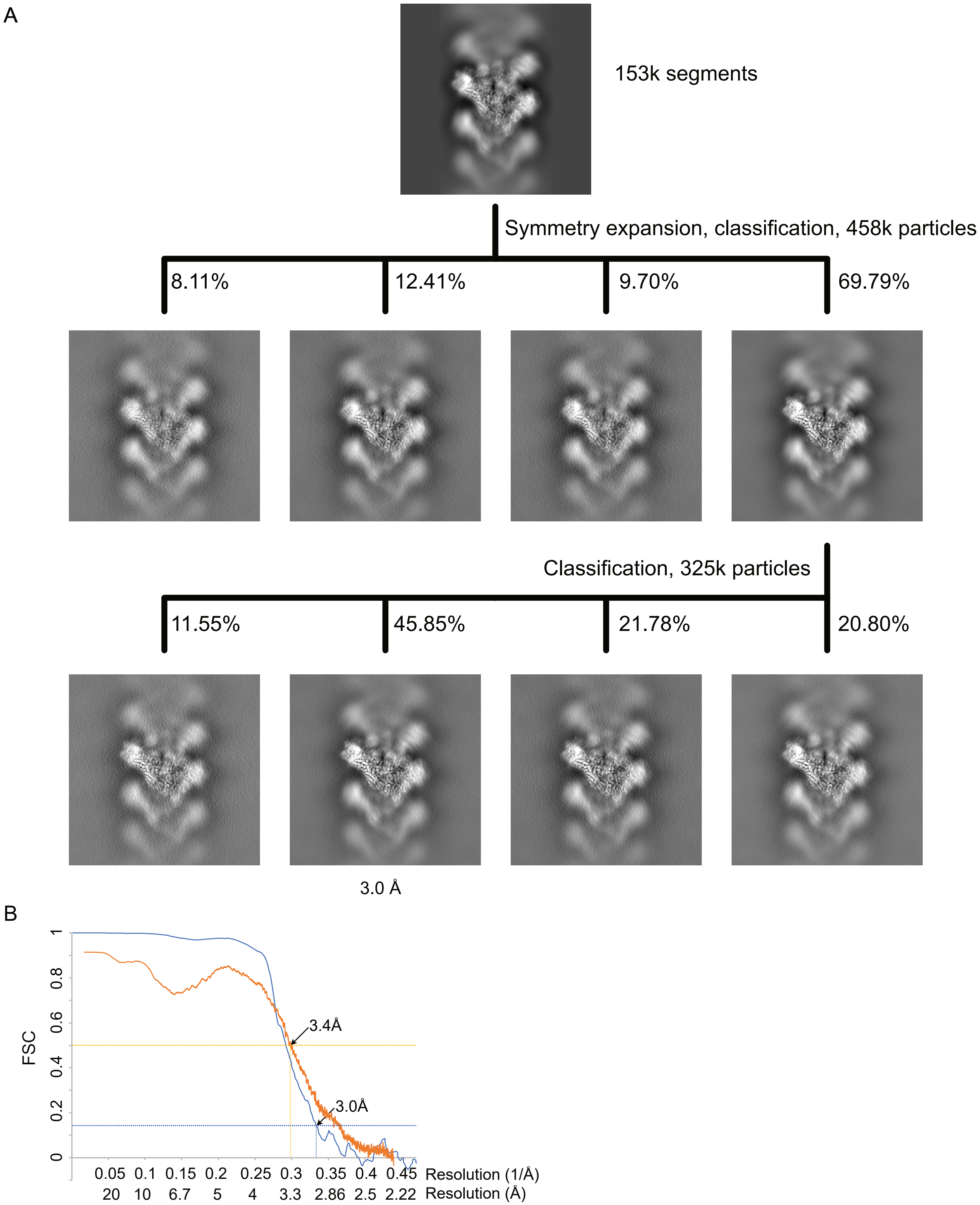


**Figure 3 - figure supplement 2: γTuRC$^{SS}$ processing and resolution**

A. Classification scheme for γTuRC$^{SS}$ processing. Images are projection images of the 3D classes obtained.
B. FSC (blue) and map to model FSC (orange) for the γTuRC$^{SS}$.

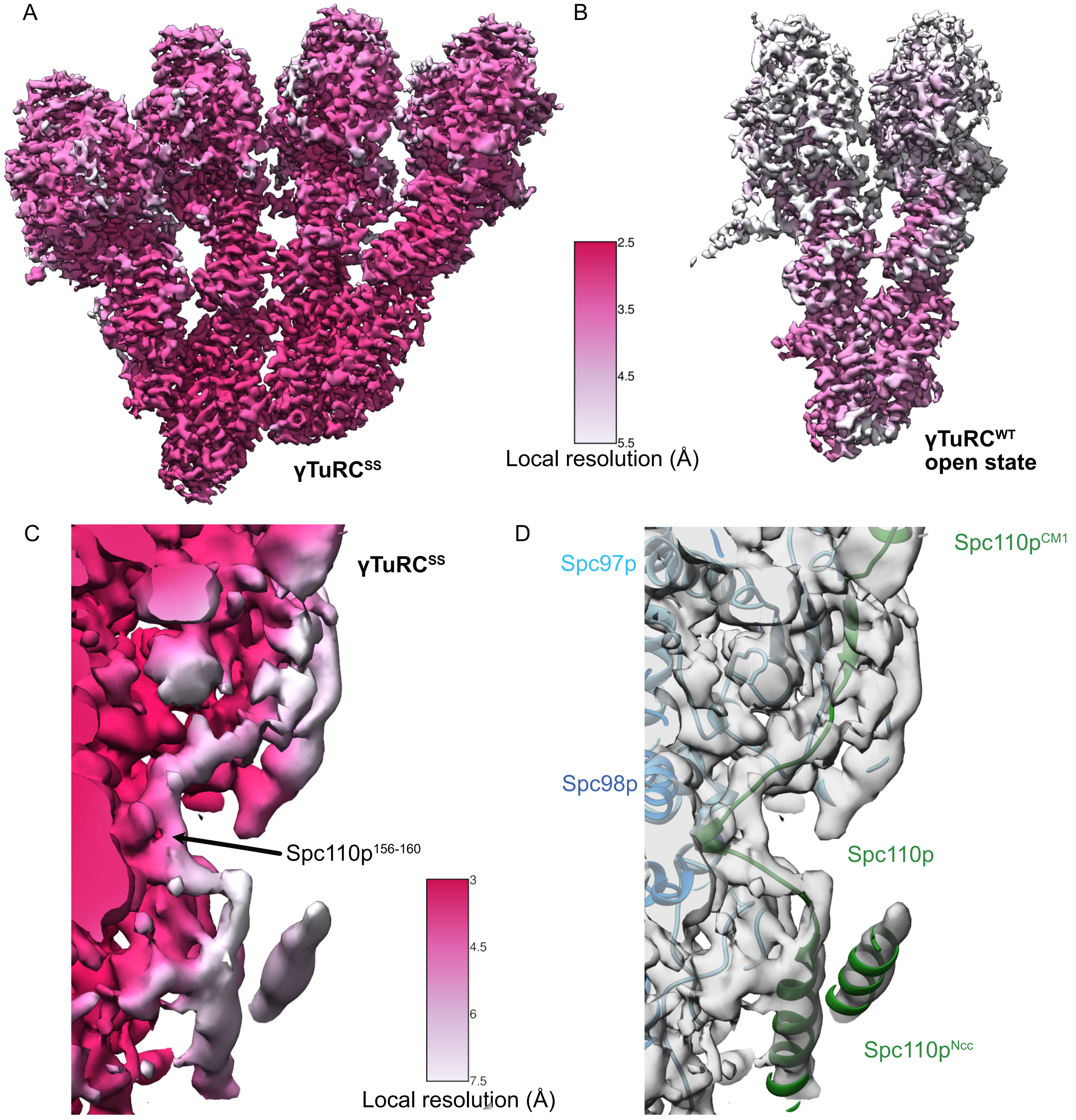


**Figure 3 - figure supplement 3: γTuRC$^{SS}$ and γTuRC$^{WT}$ local resolution maps**

A. γTuRC$^{SS}$ local resolution map. The color scale is shown between panels A and B.
B. γTuRC$^{WT}$ local resolution map. The color scale is shown between panels A and B.
C. Local resolution map highlighting the region spanning the Spc110p$^{CM1}$ and Spc110p$^{NCC}$ regions of the γTuRC$^{SS}$ reconstruction. The color scale is shown between panels C and D. The density was filtered using local resolution filtering in SPOC (https://github.com/MaximilianBeckers/SPOC; Beckers & Sachse, 2020), using the local resolutions calculated from BlocRes (See Methods). Although the density connecting the Spc110p$^{CM1}$ helix with Spc110p$^{NCC}$ is at lower resolution, a short region of higher resolution for Spc110p$^{156-160}$ and the constraints of the Spc110p$^{NCC}$ and the Spc110p$^{112-150}$ regions in building the connecting loops for Spc110p$^{161-165}$ and Spc110p$^{150-155}$ into the lower resolution density.
D. View identical to panel C showing transparent density and atomic models rendered as ribbons. Colors are as indicated by inset text.

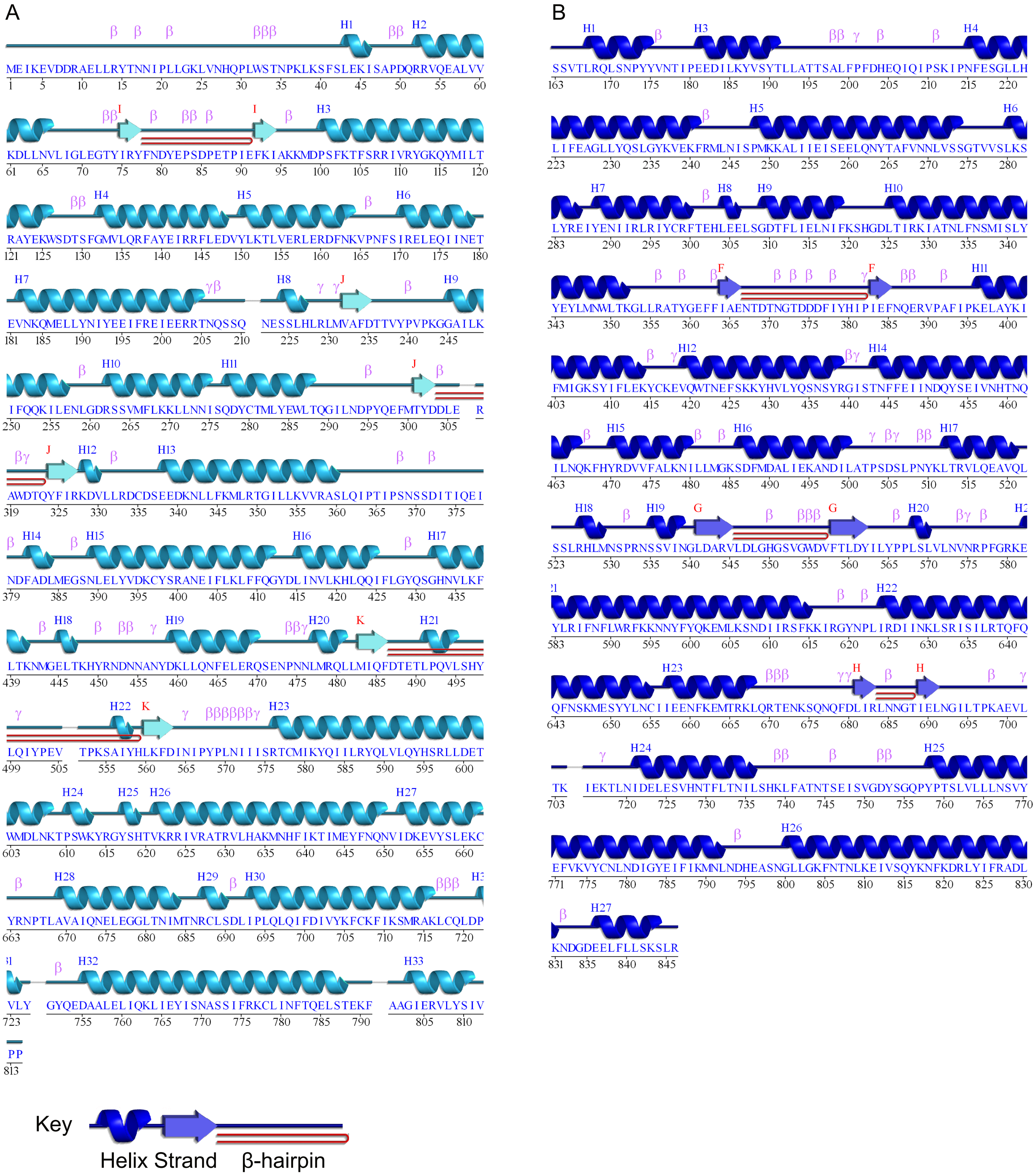


**Figure 3 - figure supplement 4: Wiring diagrams of A. Spc97p and B. Spc98p**

Helices are numbered from 1, whereas sheets are labeled from A. Turns are labeled by type. Plots were generated using the pdbsum server (Laskowski, 2009)

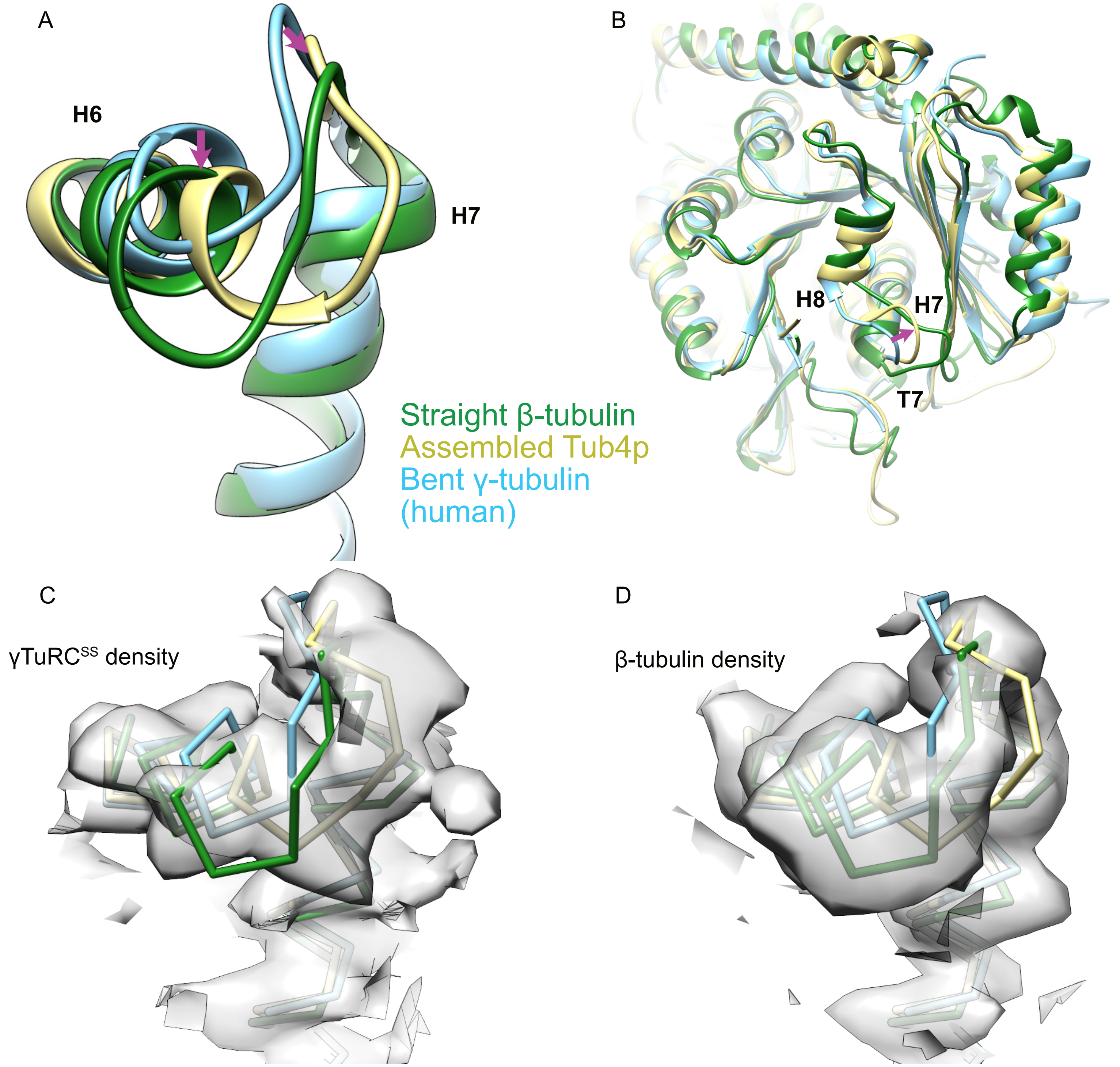


**Figure 3 - figure supplement 5: Comparison of γ-tubulin conformation between Human and Yeast γ-TuRC**

A. An alignment of γ-tubulin (γ-tubulin bound to Spc98p, khaki, this work; human free γ-tubulin, sky blue, PDB ID 1Z5W; yeast straight β-tubulin, forest green, PDB ID 5W3F) using their N-terminal domain shows that the human γ-tubulin is nearly identical to the bent crystal structure, whereas the yeast Tub4p is in an intermediate conformation more similar to the straight conformation in its H6 conformation. The magenta arrow highlights the motion of H6 and the H6-H7 loop relative to the bent conformation in both assembled and straight γ-tubulin.
B. View from the γ-tubulin:spc98p interface of a global overlay of γ-tubulin bound to Spc98p and the human free γ-tubulin structure shows a motion of T7 loop at the γ-tubulin:Spc98p interface during assembly. The magenta arrow highlights the motion of the T7 loop relative to the bent conformation in both assembled and straight γ-tubulin.
C. View of aligned yeast γ-tubulin, human free γ-tubulin and yeast straight β-tubulin, colored and aligned as in panel A, showing that the density from the yeast γ-tubulin as seen in the γTuRC$^{SS}$ reconstruction is clearly distinct from the bent and straight conformations. Density is filtered at 4.2 Å for clarity.
D. View of aligned yeast γ-tubulin, human free γ-tubulin and yeast straight β-tubulin, colored and aligned as in panel A, showing that the density from the yeast β-tubulin filament reconstruction is clearly distinct from the γTuRC$^{SS}$ and bent conformations.
Density in panels C and D was zoned within 2.8 Å of their respective models.

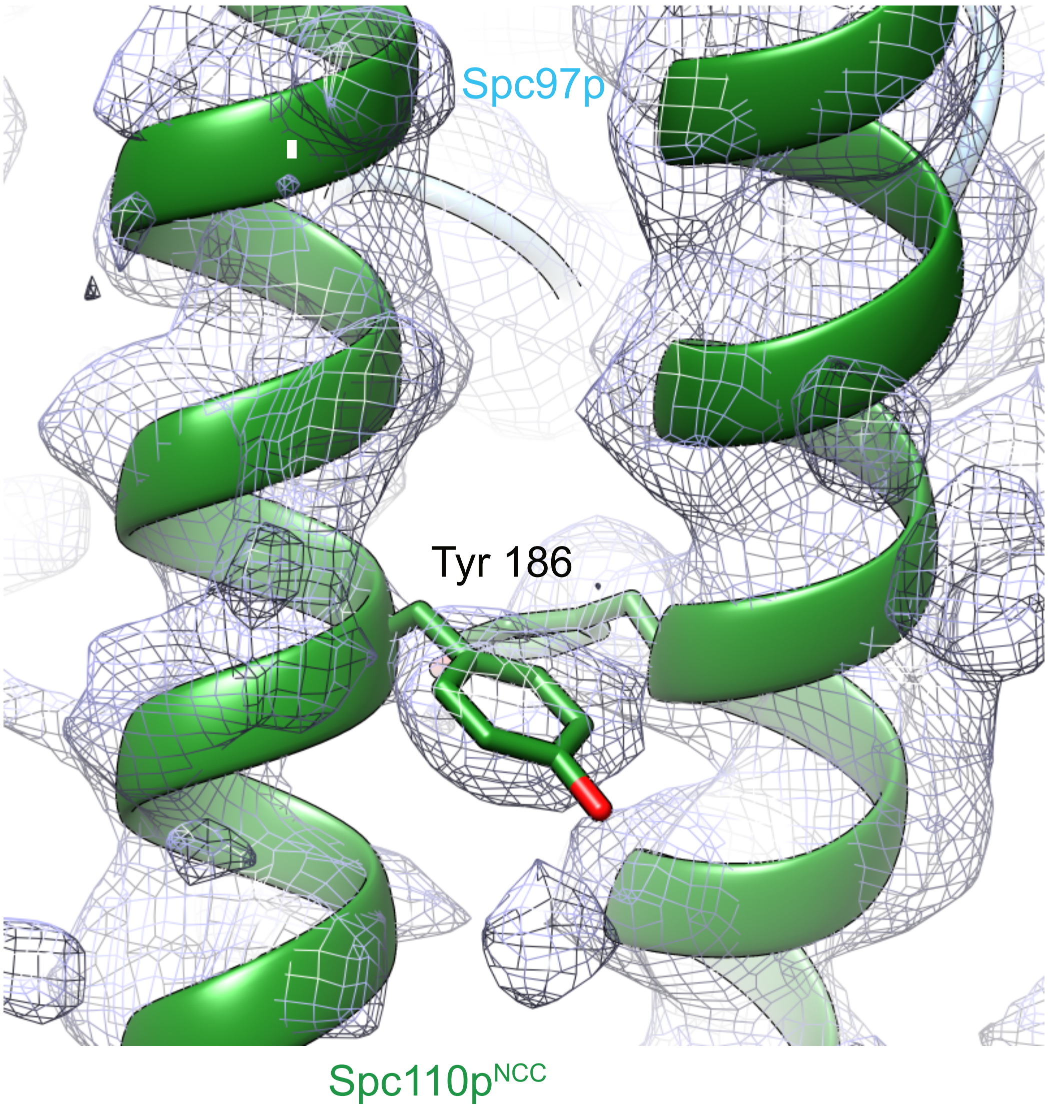


**Figure 4 - figure supplement 1: Spc110p$^{NCC}$ structure (forest green) near the tyrosine 186 side chain fitted into ~4.2Å low pass filtered density from the γTuRC$^{SS}$ reconstruction.**

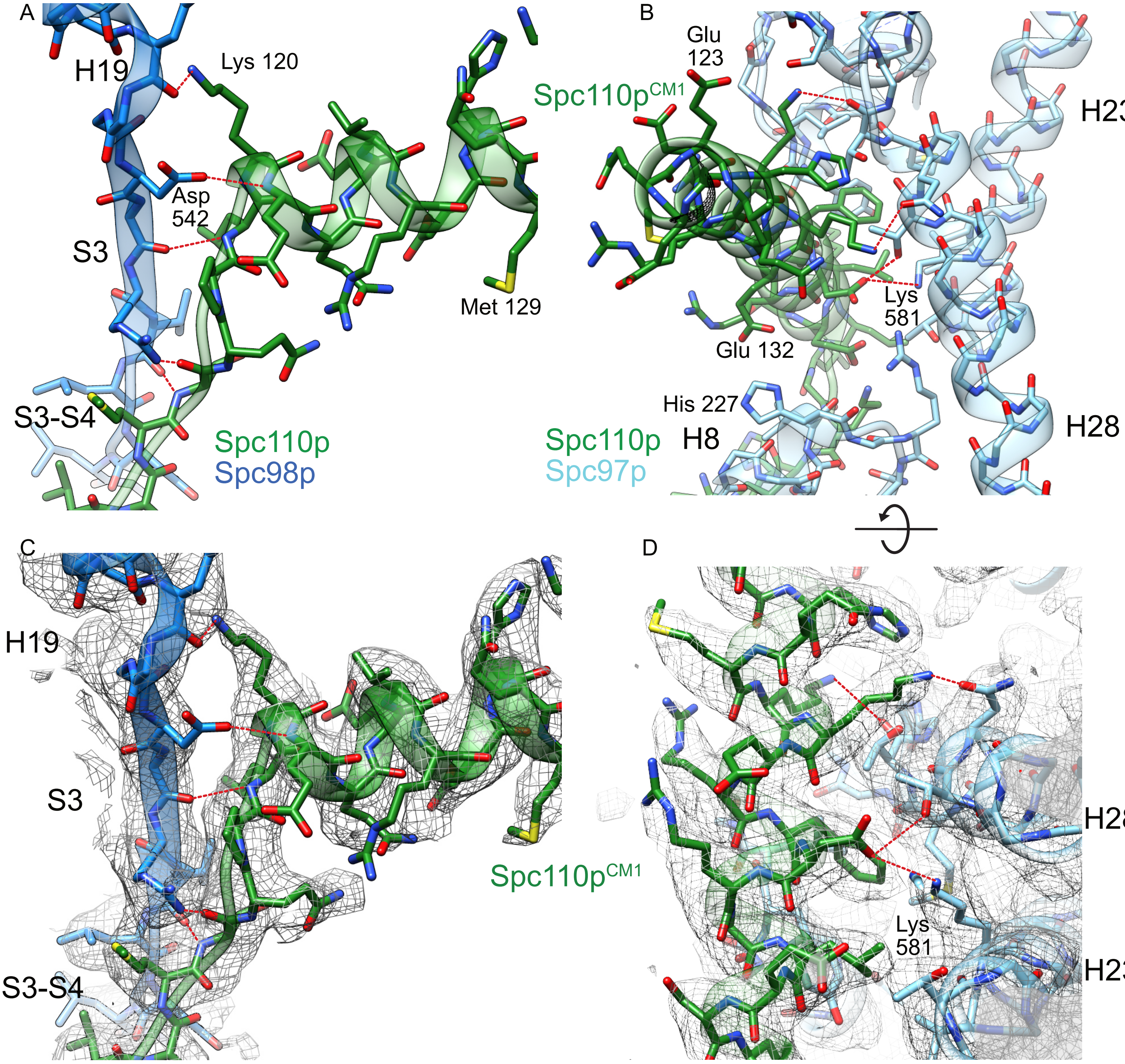


**Figure 4 - figure supplement 2: Helix dipole interactions define the CM1 binding site on Spc98p**

A. View of the Spc110p$^{CM1}$ motif binding site with Spc98p with the H19-S3 region, colored as in Fig 4. Spc110p$^{CM1}$:Spc98p hydrogen bond interactions are indicated in red. All of the hydrogen bonds between Spc110p and Spc98p have a backbone atom as one of the hydrogen bonds partners.
B. View of the CM1 binding site with Spc97p. Side chains interacting with the Spc110p$^{CM1}$ helix are shown. Spc110p$^{CM1}$:Spc97p hydrogen bond interactions are indicated in red.
C. View of the Spc110p$^{CM1}$ motif binding site with Spc98p with the H19-S3 region, with the model overlaid with density. Density in this panel is filtered at 3.0 Å, and zoned within 2.6 Å from the model.
D. View of the CM1 binding site with Spc97p, with the model overlaid with density. The view has Density in this panel is filtered at 3.5 Å, and density is unmasked.
Hydrogen bonds labeled in panels A and B were identified using the "FindHBond" GUI as implemented in Chimera, using the default parameters for relaxing H-bond constraints of 0.4 Å and 20.0 degrees.

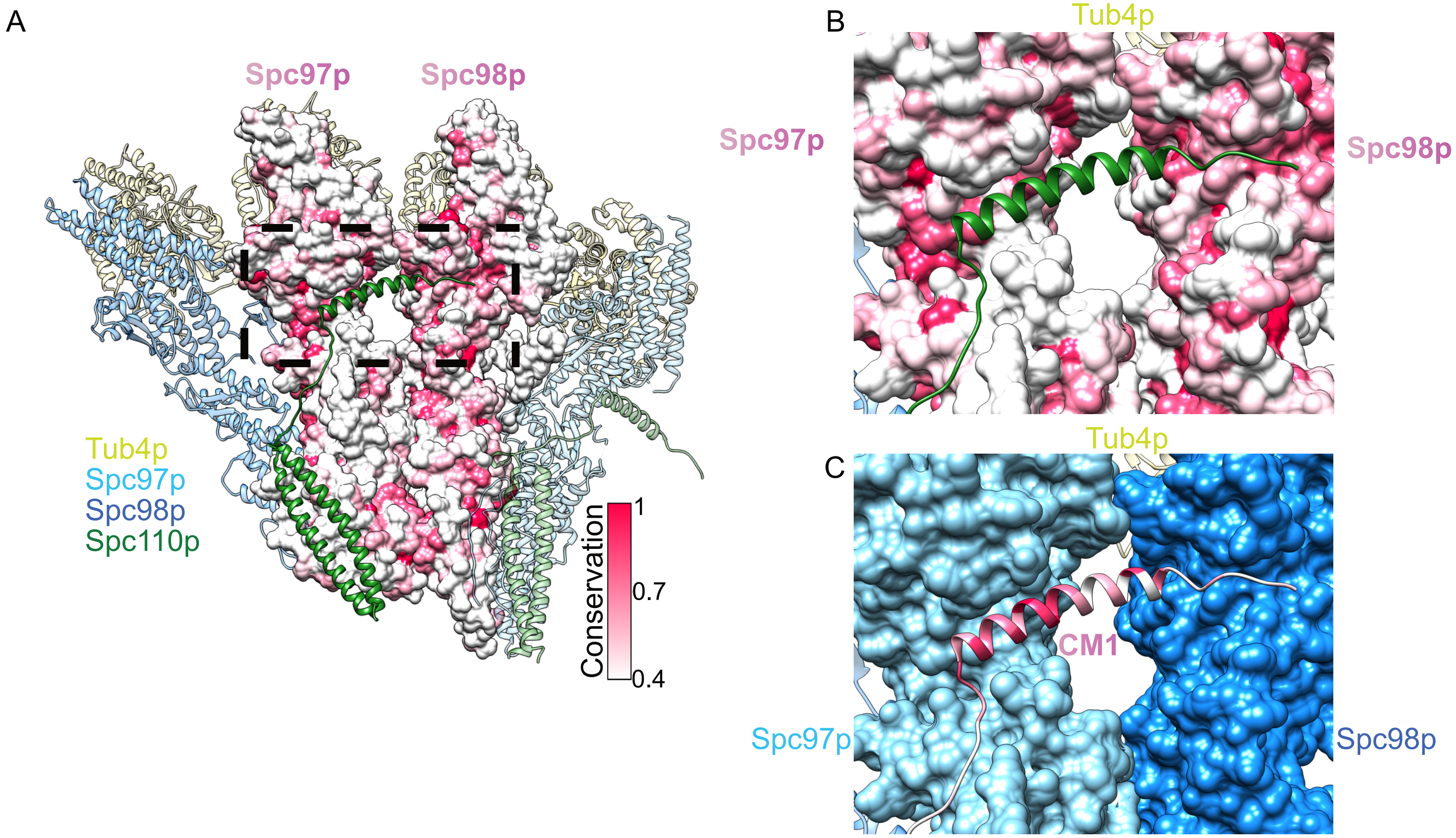


**Figure 4 - figure supplement 3: Conserved binding interface with the CM1 motif**

A. Overview of a dimer of γTuSC colored as in Figure 2, with the central Spc97p/98p colored according to their conservation.
B. View of the Spc110p$^{CM1}$ binding site at the Spc97p/98p C-terminus, with Spc97p/98p colored according to their conservation, as in panel A.
C. View of the Spc110p$^{CM1}$ binding site at the Spc97p/98p C-terminus, with Spc110p colored according to its conservation, with the same scale as panel A. Spc97p/98p are colored as in the figure legend.

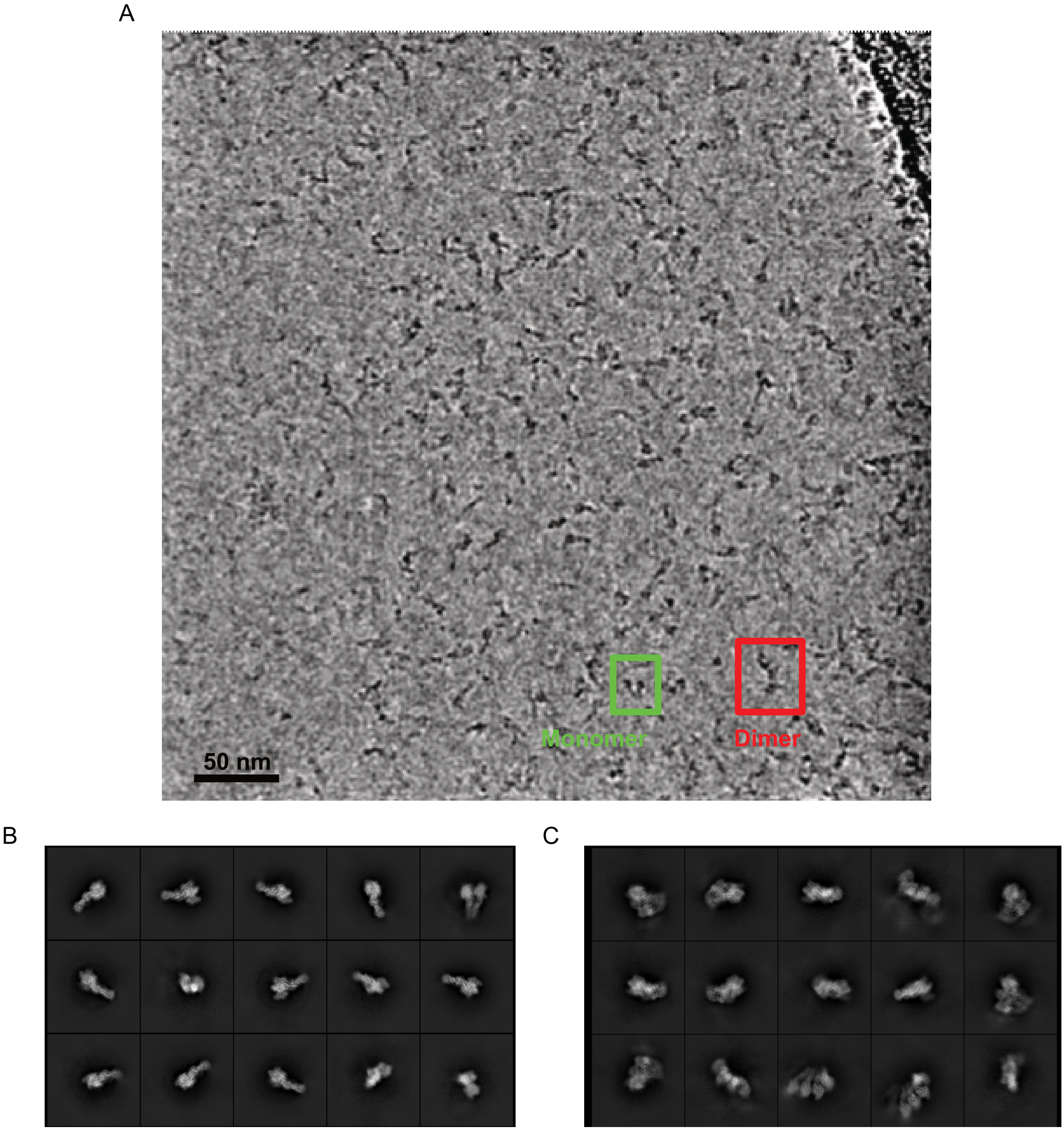


**Figure 5 - figure supplement 1: A mixture of compositional states is observed**

A. A filtered micrograph showing raw particles shows well dispersed single particles and a mixture of γ-TuSC monomers and dimers.
B. Representative examples of classes obtained by unsupervised classification of monomers
C. Representative examples of classes obtained by unsupervised classification of dimers.

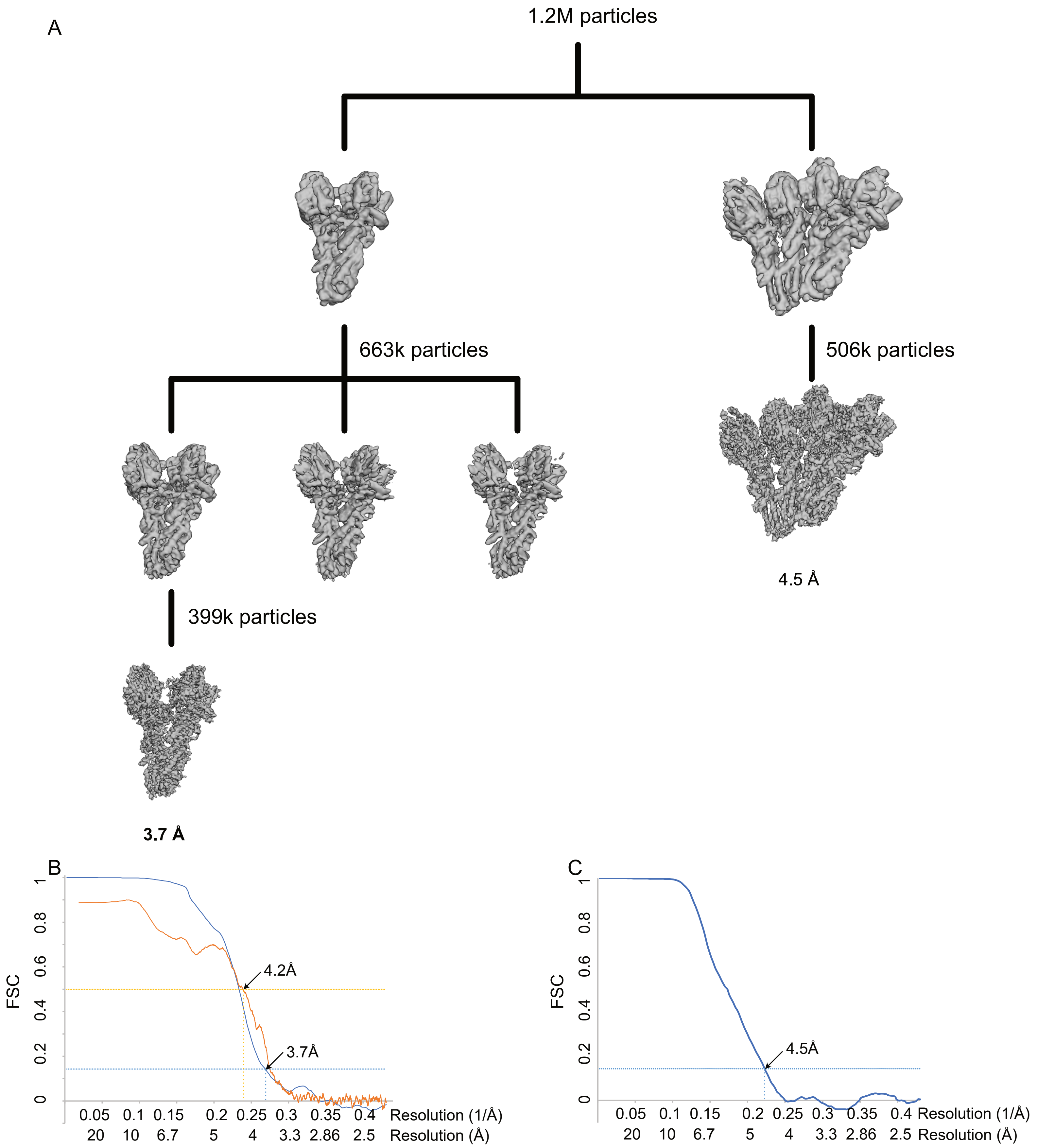


**Figure 5 - figure supplement 2: WT γTuSC processing and resolution**

A. Classification scheme for WT γTuSC monomer processing
B. Classification scheme for WT γTuSC dimer processing
C. FSC (blue) and map to model FSC for the closed WT γTuSC monomer
D. FSC (blue) and map to model FSC for the open WT γTuSC dimer

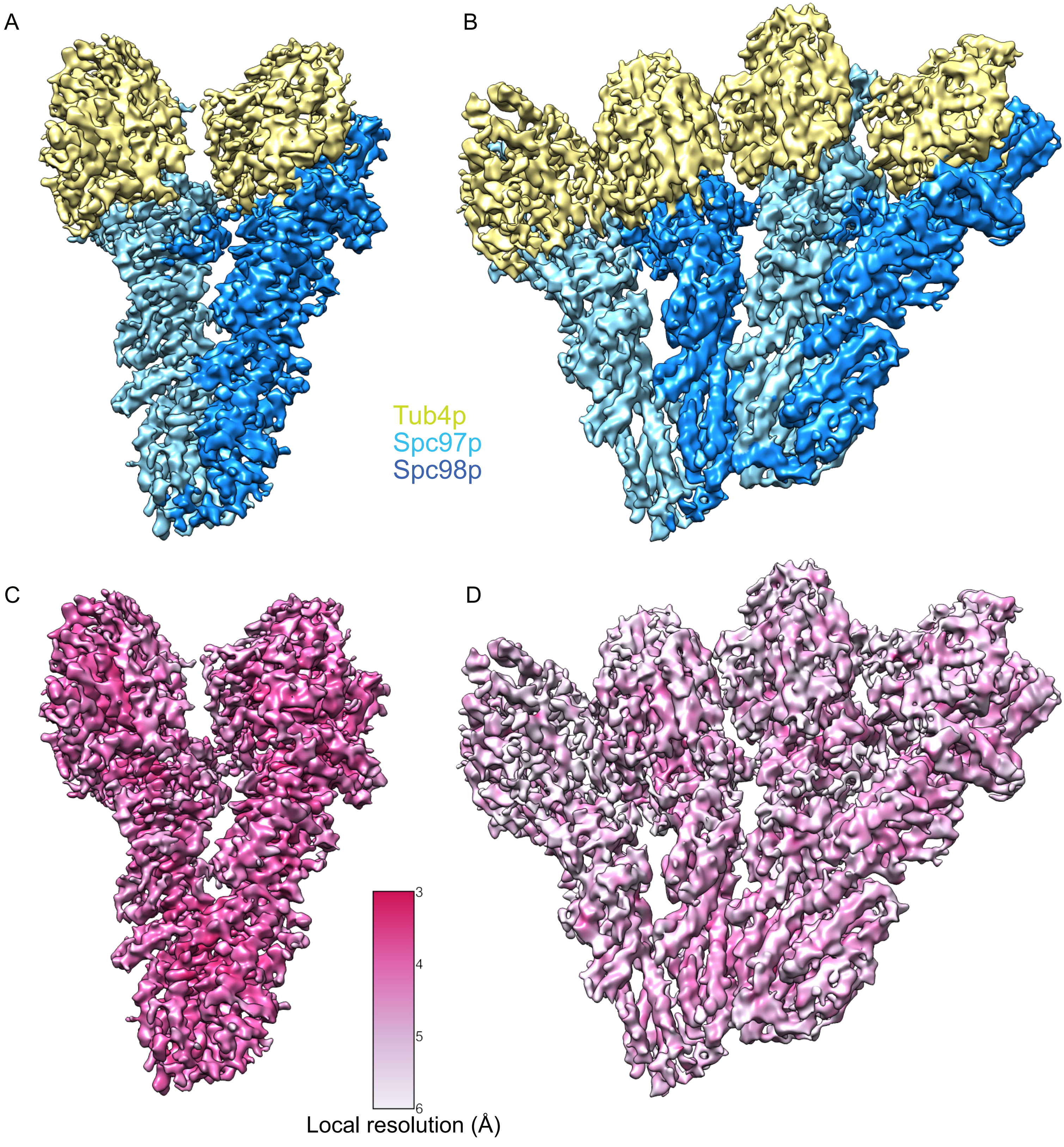


**Figure 5 - figure supplement 3: Segmented single-particle reconstructions of γTuSC monomer and dimer**

A. ~3.7 Å reconstruction γTuSC showing Spc97p (sky blue), Spc98p (dodger blue) and γ-tubulin (khaki)
B. ~4.5 Å reconstruction of a dimer of γTuSC colored as in panel A.
C. ~3.7 Å reconstruction γTuSC colored according to its local resolution.
D. ~4.5 Å reconstruction of a dimer of γTuSC colored according to its local resolution.
Color scale for panels C and D is shown in figure inset.
Disconnected density smaller than 5 Å was hidden using the "Hide Dust" command in Chimera.

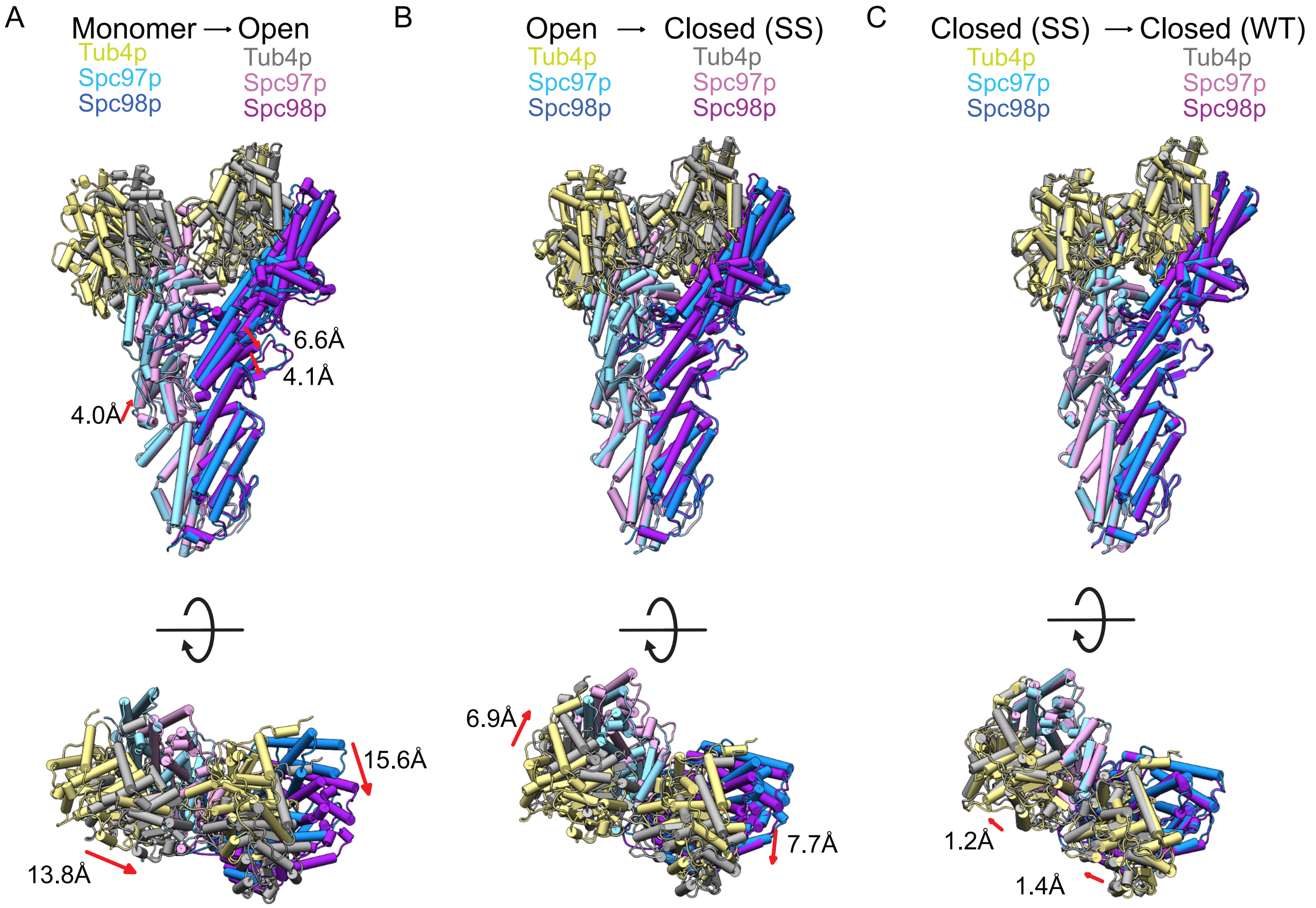


**Figure 5 - figure supplement 4: Conformational changes in γTuSC during assembly and activation**

A. Views of the monomeric γTuSC aligned with an open γTuRC$^{WT}$ filament monomer using the N-terminal helical bundles of Spc97/98p. Monomeric γTuSC is colored as in Fig. 2, and the γTuSC$^{WT}$ filament has Spc97p in light pink, Spc98p in purple, and Tub4p colored in gray.
B. Views of a γTuRC$^{WT}$ filament monomer aligned with a γTuRC$^{ss}$ filament monomer using the N-terminal helical bundles of Spc97/98p. γTuSC$^{WT}$ is colored as in Fig. 2, and γTuSC$^{ss}$ has Spc97p in light pink, Spc98p in purple, and Tub4p colored in gray.
C. Views of a γTuSC$^{ss}$ filament monomer aligned with a γTuSC$^{WT}$ closed filament monomer using the N-terminal helical bundles of Spc97/98p. γTuSC$^{ss}$ is colored as in Fig. 2, and γTuSC$^{WT}$ has Spc97p in light pink, Spc98p in purple, and Tub4p colored in gray.

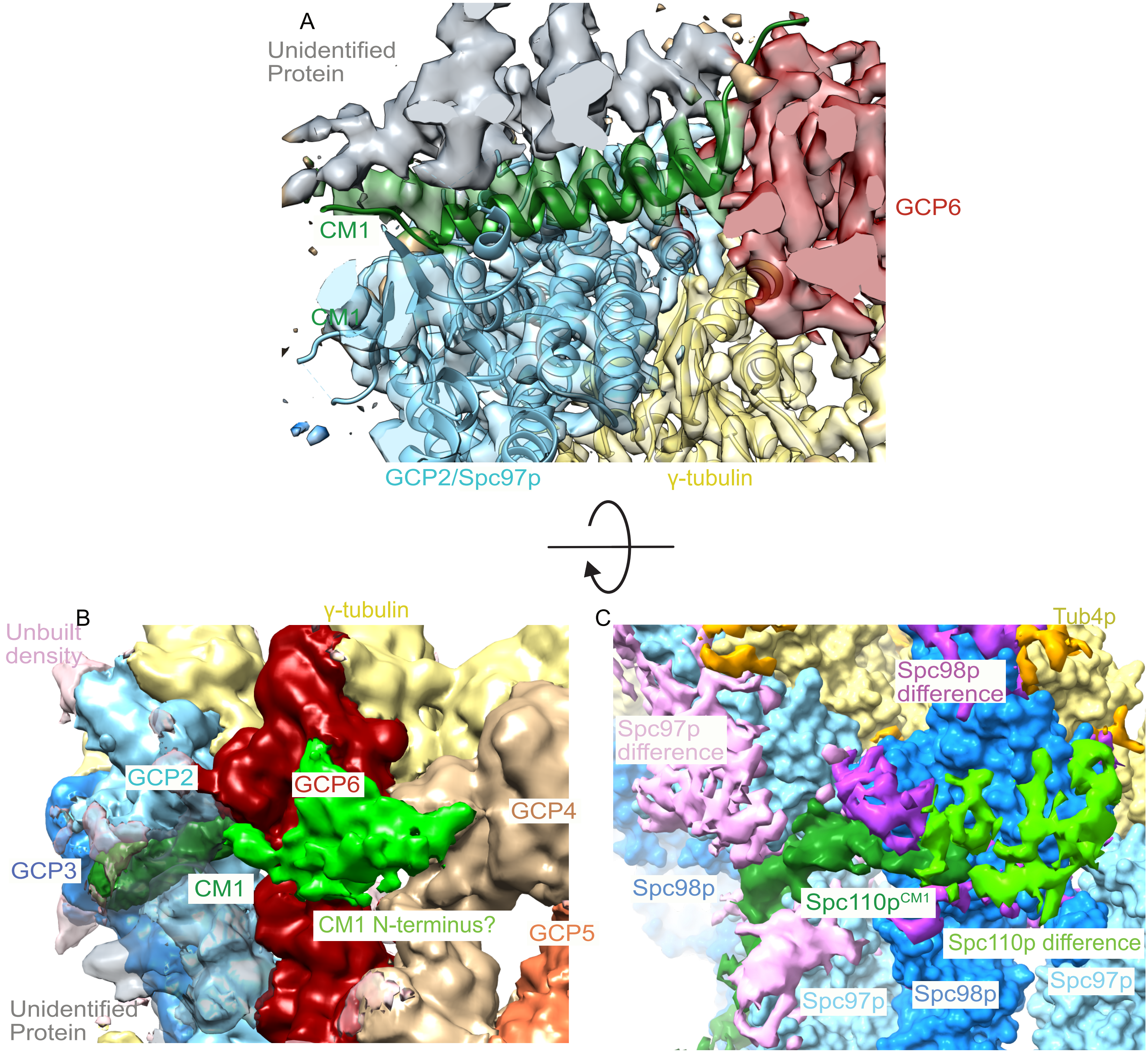


**Figure 7 - figure supplement 1: A CM1 helix binds between GCP2 and GCP6 in human γ-TuRC**

A. Human γ-TuRC density (EMDB ID 21068) is fitted with the C-terminal region of Yeast GCP2, γ-Tubulin and CM1 helix (this work), showing that a CM1 helix binds between GCP2 and GCP6 in human γ-TuRC.
B. Low resolution filtered human γ-TuRC density (EMDB ID 21068) density shows a similar N-terminal density extending from the CM1 helix towards the adjacent GCP as observed in the (C) yeast difference map (this work).

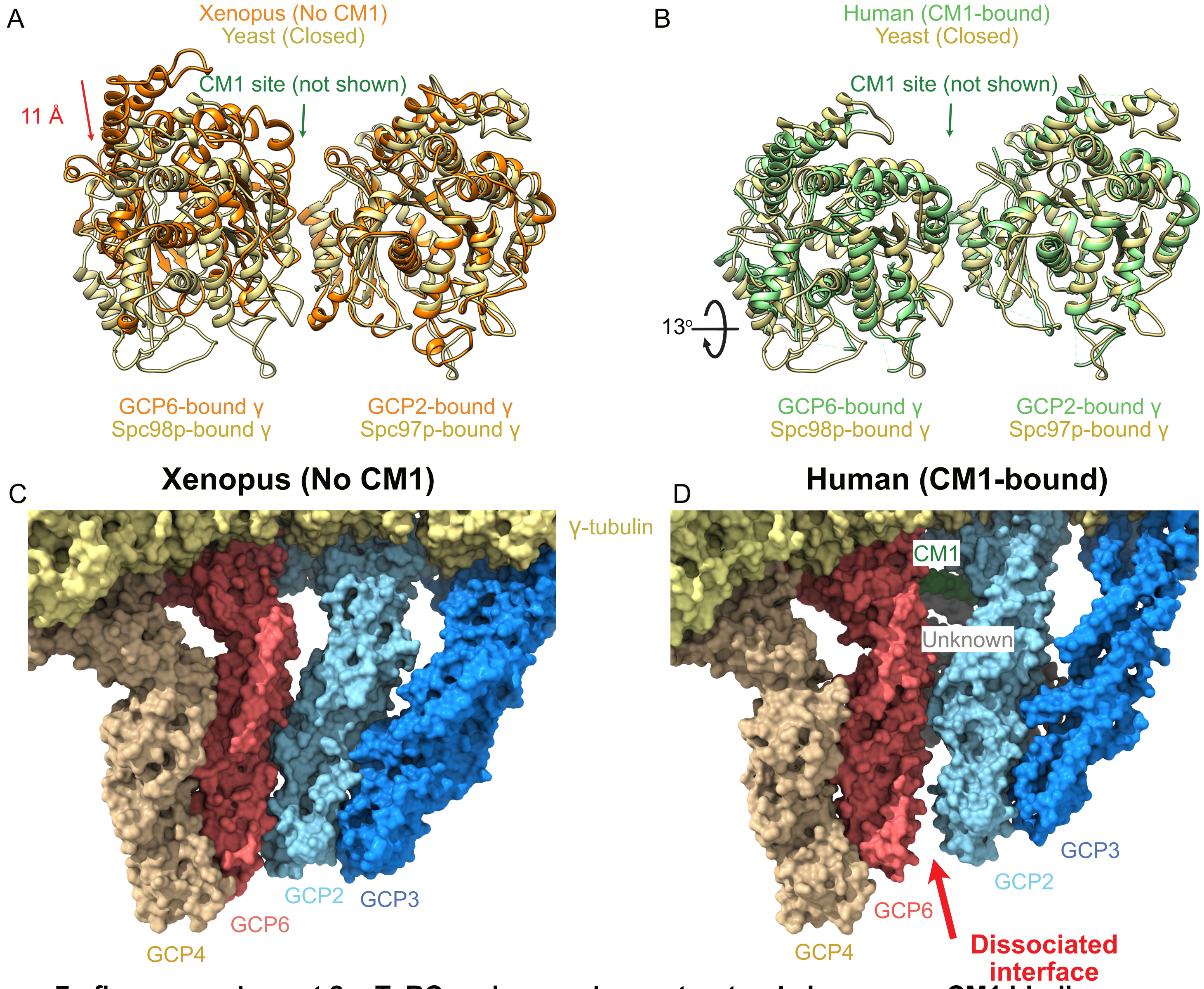


**Figure 7 - figure supplement 2: γTuRC undergoes large structural changes on CM1 binding**

A-B. Overlay of the two γ-tubulins adjacent to the CM1 binding site observed in the human γTuRC structure for **(A)** xenopus (No CM1 bound) and **(B)** human (CM1-bound) highlighting the large motion upon CM1 binding. Alignment is based on the GCP2-bound γ-tubulin.
C-D. Surface representation of the GCP2/6 interface shows a well formed interface in the xenopus (no CM1 bound) γ-TuRC structure, in contrast to the dissociated interface observed in the human (CM1-bound) γ-TuRC structure.
