## Supplemental Movie Legends for "CM1-driven assembly and activation of Yeast γ-Tubulin Small Complex underlies microtubule nucleation"

**Movie S1. Conformational Changes of γTuSC During Assembly**

Morph of the monomeric γTuSC conformation to the conformation adopted by a $\gamma TuRC^{WT}$ filament monomer in the open state. The γTuSC is colored with Tub4p in khaki, Spc97p in light blue and Spc98p in light blue (as in Fig. 2). Morphs were calculated in Chimera (Pettersen et al., 2004) using a linear interpolation method. Corresponds to transition shown in Fig. 5 – figure supplement 4A.

**Movie S2. Conformational Changes of γTuRC During Activation**

Morph of a $\gamma TuRC^{WT}$ filament monomer in the open conformation to the conformation adopted by a $\gamma TuRC^{SS}$ filament monomer in the closed state. The γTuSC is colored with Tub4p in khaki, Spc97p in light blue and Spc98p in light blue (as in Fig. 2). Morphs were calculated in Chimera (Pettersen et al., 2004) using a linear interpolation method. Corresponds to transition shown in Fig. 5 – figure supplement 4B.

**Movie S3. Morph of $\gamma TuRC^{SS}$ and $\gamma TuRC^{WT}$ closed states shows minimal changes**

Morph of a $\gamma TuRC^{SS}$ filament monomer in the closed state to the conformation adopted by a $\gamma TuRC^{WT}$ filament monomer in the closed state. The γTuSC is colored with Tub4p in khaki, Spc97p in light blue and Spc98p in light blue (as in Fig. 2). Morphs were calculated in Chimera (Pettersen et al., 2004) using a linear interpolation method. Corresponds to transition shown in Fig. 5 – figure supplement 4C.
