## Supplementary material for "CM1-driven assembly and activation of Yeast γ-Tubulin Small Complex underlies microtubule nucleation": Computational Methods Supplement

***Supplementary Computational Methods***

#### *Localizing Spc110 on γTuSC using integrative structure determination*

The localization of Spc110 on γTuSC using integrative structure determination proceeded through four stages (Fig. 2 – figure supplement 3) (Alber et al., 2007; Rout & Sali, 2019; Russel et al., 2012): (1) gathering data, (2) representing the system and translating data into spatial restraints, (3) structural sampling to produce an ensemble of structures that satisfies the restraints, and (4) analyzing and validating the ensemble structures and data. The modeling protocol (*i.e*., stages 2, 3, and 4) was scripted using the *Python Modeling Interface* (PMI) package, a library for modeling macromolecular complexes based on our open-source *Integrative Modeling Platform* (IMP) package, version 2.8 (https://integrativemodeling.org) (Russel *et al.*, 2012). The current procedure is an updated version of previously described protocols (Kim et al., 2018; Viswanath, Bonomi, et al., 2017; Wang et al., 2017; Webb et al., 2018). Files containing the input data, scripts, and output results are available at https://salilab.org/gtuscSpc110 as well as the nascent Protein Data Bank archive for integrative structures (https://pdb-dev.wwpdb.org/) with deposition codes PDBDEV_00000077 (γTuSC monomer with Spc110p monomer) PDBDEV_00000078 (γTuSC monomer with Spc110p dimer) PDBDEV_00000079 (γTuSC dimer with two Spc110p dimers).

#### Stage 1: Gathering data

Chemical crosslinks between between Spc110 and γTuSC were identified by mass spectrometry of samples containing either Spc110$^{1-220}$-GCN4 dimer or Spc110$^{1-401}$-GST, informing the localization of Spc110 relative to γTuSC (Fig. 2 – figure supplement 1). γTuSC structure used was obtained from the PDB (code 5FLZ); it was determined primarily based on a cryo-EM density map of the disulfide-stabilized γTuSC filament at 6.9 Å resolution (EMDB code: 2799) (Greenberg et al., 2016; Kollman et al., 2015). Representation of Spc110$^{1-220}$ relied on (i) crystal structure of Spc110 N-terminal coiled-coil (NCC) domain (Fig 2B) and (ii) failure to detect related sequences of known structure in the rest of the Spc110 sequence by HHPred (Söding, Biegert, & Lupas, 2005).

#### Stage 2: Representing the system and translating data into spatial restraints

Information about the modeled system (above) can in general be used for defining its representation, defining the scoring function that guides sampling of alternative models, limiting sampling, filtering of good-scoring models obtained by sampling, and final validation of the models (Fig. 2 – figure supplement 3). Here, the "flexible" representation for most of Spc110$^{1-220}$ reflects the absence of known related structures. The γTuSC and Spc110 NCC domain representations rely on their atomic structures. The scoring function relies on chemical cross-links, excluded volume, and sequence connectivity.

An optimal representation facilitates accurate formulation of spatial restraints as well as efficient and complete sampling of good-scoring solutions, while retaining sufficient detail without overfitting, so that the resulting models are maximally useful for subsequent biological analysis (Viswanath & Sali, 2019). We first used a representation where a single γTuSC was bound to an $Spc110^{1-220}$ dimer. To maximize computational efficiency while avoiding using too coarse a representation, we represented the system in a multi-scale fashion. A rigid body consisting of multiple beads was defined for γTuSC and the Spc110 NCC ($Spc110^{164-203}$). In a rigid body, the beads have their relative distances constrained during conformational sampling, whereas in a flexible string the beads are restrained by the scoring function (below). Rigid bodies were coarse-grained using one-residue beads, whose coordinates were those of the corresponding $C_\alpha$ atoms. The remaining regions in γTuSC without an atomic model were represented by a flexible string of beads encompassing 20 residues each. Due to lack of acceptable comparative models, and knowing that a large region of the N-terminus of Spc110 lacks secondary structure (Fig. 2A), we used a flexible string of 5-residue beads each to represent regions of $Spc110^{1-220}$ other than the coiled-coil domains. Additionally, we modeled a single γTuSC bound to an $Spc110^{1-220}$ monomer, as well as a complex of two adjacent γTuSCs each bound to an $Spc110^{1-220}$ dimer.

With this representation in hand, we next encoded the spatial restraints into a scoring function based on the information gathered in Stage 1, as follows:

(1) *Cross-link restraints:* The cross-links (Fig. 2 – figure supplement 1) were used to construct the Bayesian scoring function (Rieping, Habeck, & Nilges, 2005) that restrained the distances spanned by the cross-linked residues (Shi et al., 2014).

(2) *Excluded volume restraints*: The excluded volume restraints were applied to each bead, using the statistical relationship between the volume and the number of residues that it covered (Alber et al., 2007).

(3) *Sequence connectivity restraints*: We applied the sequence connectivity restraints, using a harmonic upper distance bound on the distance between consecutive beads in a subunit, with a threshold distance equal to twice the sum of the radii of the two connected beads. The bead radius was calculated from the excluded volume of the corresponding bead, assuming standard protein density (Alber et al., 2007; Shi et al., 2014).

**Stage 3: Structural sampling to produce an ensemble of structures that satisfies the restraints**

We aimed to maximize the precision at which the sampling of good-scoring solutions was exhaustive (Stage 4). We sampled the positions of flexible Spc110 beads and the flexible linkers of γTuSC. The search for good-scoring models relied on Gibbs sampling, based on the Metropolis Monte Carlo algorithm (Wang et al., 2017). The positions of the γTuSC rigid body and the Spc110 NCC rigid body were fixed, while the initial positions of flexible γTuSC and Spc110 beads were randomized. The Monte Carlo moves included random translations of individual beads in the flexible segments of γTuSC and Spc110 (up to 3 Å). A model was saved every 10 Gibbs sampling steps, each consisting of a cycle of Monte Carlo steps that moved every moving bead once.

This sampling produced a total of 30 million models from 50 independent runs, requiring ~2 days on 200 CPU cores. For the most detailed specification of the sampling procedure, see the IMP modeling script (https://salilab.org/gtuscSpc110). We only consider for further analysis the ~3,000 good-scoring models that satisfy the input datasets within their uncertainties (below).

**Stage 4: Analyzing and validating the ensemble structures and data**

Input information and output structures need to be analyzed to estimate structure precision and accuracy, detect inconsistent and missing information, and to suggest more informative future experiments. We used the analysis and validation protocol published earlier (Alber et al., 2007; Kim et al., 2018; Rout & Sali, 2019; Viswanath, Bonomi, et al., 2017; Viswanath, Chemmama, Cimermancic, & Sali, 2017): Assessment began with the clustering of the models and estimating their precision based on the variability in the ensemble of good-scoring structures, and quantification of the structure fit to the input information. These validations are based on the nascent wwPDB effort on archival, validation, and dissemination of integrative structure models (Burley et al., 2017; Sali et al., 2015). We now discuss each one of these points in turn.

*(1) Clustering and structure precision*

An ensemble of good-scoring structures needs to be analyzed in terms of the precision of its structural features (Viswanath, Chemmama, et al., 2017). The precision of a component position can be quantified by its variation in an ensemble of superposed good-scoring structures. It can also be visualized by the localization probability density for each of the components of the model.

As described above, integrative structure determination of the γTuSC-Spc110$^{1\text{-}220}$ dimer complex resulted in effectively a single good-scoring solution, at the precision of 23.3 Å (Fig. 2 – figure supplement 4). The precision is the bead RMSD from the cluster centroid model averaged over all models in the cluster. Additionally, the sampling precisions for the γTuSC-Spc110$^{1\text{-}220}$ monomer complex and complex of two adjacent γTuSC-Spc110$^{1\text{-}220}$ dimers were 13.4 Å and 34.1 Å respectively.

*(2) Fit to input information*

An accurate structure needs to satisfy the input information used to compute it. The cluster of models of the γTuSC-Spc110$^{1\text{-}220}$ dimer complex satisfied 90.9% (92.8%) of the EDC (DSS) crosslinks; a crosslink is satisfied by a cluster of models if the corresponding Cα-Cα distance in any model in the cluster is less than 35 Å (25 Å) for DSS (EDC) crosslinks. Additionally, the γTuSC-Spc110$^{1\text{-}220}$ monomer complex satisfied 81.8% (80.9%) of EDC (DSS) crosslinks and the complex of two adjacent γTuSC-Spc110$^{1\text{-}220}$ dimers satisfied 100% (94%) of EDC (DSS) crosslinks. The remainder of the restraints are harmonic, with a specified standard deviation. The cluster generally satisfied at least 95% of restraints of each type (excluded volume and sequence connectivity). A restraint is satisfied by a cluster of models if the restrained distance in any model in the cluster (considering restraint ambiguity) is violated by less than 3 standard deviations, specified for the restraint. Most of the violations are small, and can be rationalized by local structural fluctuations, coarse-grained representation of the model, and/or finite structural sampling.

### ***Supplemental References***


Alber, F., Dokudovskaya, S., Veenhoff, L., Zhang, W., Kipper, J., Devos, D., . . . Sali, A. (2007). Determining the architectures of macromolecular assemblies. *Nature, 450*(7170), 683-694.

Burley, S. K., Kurisu, G., Markley, J. L., Nakamura, H., Velankar, S., Berman, H. M., . . . Trewhella, J. (2017). PDB-Dev: A Prototype System for Depositing Integrative/Hybrid Structural Models. *Structure, 25*(9), 1317-1318

Greenberg, C. H., Kollman, J., Zelter, A., Johnson, R., MacCoss, M. J., Davis, T. N., . . . Sali, A. (2016). Structure of γ-tubulin small complex based on a cryo-EM map, chemical cross-links, and a remotely related structure. *J Struct Biol, 194*(3), 303-310

Kim, S. J., Fernandez-Martinez, J., Nudelman, I., Shi, Y., Zhang, W., Raveh, B., . . . Rout, M. P. (2018). Integrative Structure and Functional Anatomy of a Nuclear Pore Complex. *Nature, 555*(7697), 475-482

Kollman, J., Greenberg, C., Li, S., Moritz, M., Zelter, A., Fong, K., . . . Agard, D. (2015). Ring closure activates yeast γTuRC for species-specific microtubule nucleation. *Nat Struct Mol Biol, 22*, 132-137.

Rieping, W., Habeck, M., & Nilges, M. (2005). Inferential structure determination. *Science, 309*(5732), 303-306.

Rout, M. P., & Sali, A. (2019). Principles for Integrative Structural Biology Studies. *Cell, 177*(6), 1384-1403. doi:10.1016/j.cell.2019.05.016

Russel, D., Lasker, K., Webb, B., Velazquez-Muriel, J., Tjioe, E., Schneidman-Duhovny, D., . . . Sali, A. (2012). Putting the pieces together: integrative structure determination of macromolecular assemblies. *PLoS Biol, 10*(1), e1001244.

Sali, A., Berman, H., Schwede, T., Trewhella, J., Kleywegt, G., Burley, S., . . . Westbrook, J. (2015). Outcome of the First wwPDB Hybrid/Integrative Methods Task Force Workshop. *Structure, 23*(7), 1156-1167.

Shi, Y., Fernandez-Martinez, J., Tjioe, E., Pellarin, R., Kim, S. J., Williams, R., . . . Chait, B. (2014). Structural characterization by cross-linking reveals the detailed architecture of a coatomer-related heptameric module from the nuclear pore complex. *Mol Cell Proteomics, 13*, 2927-2943.

Söding, J., Biegert, A., & Lupas, A. N. (2005). The HHpred interactive server for protein homology detection and structure prediction. *Nucleic acids research, 33*(suppl_2), W244-W248.

Viswanath, S., Bonomi, M., Kim, S. J., Klenchin, V. A., Taylor, K. C., Yabut, K. C., . . . Muller, E. G. (2017). The molecular architecture of the yeast spindle pole body core determined by Bayesian integrative modeling. *Mol Biol Cell, 28*(23), 3298-3314.

Viswanath, S., Chemmama, I. E., Cimermancic, P., & Sali, A. (2017). Assessing Exhaustiveness of Stochastic Sampling for Integrative Modeling of Macromolecular Structures. *Biophys J, 113*(11), 2344-2353. doi:10.1016/j.bpj.2017.10.005

Viswanath, S., & Sali, A. (2019). Optimizing model representation for integrative structure determination of macromolecular assemblies. *Proc Natl Acad Sci U S A, 116*(2), 540-545. doi:10.1073/pnas.1814649116

Wang, X., Chemmama, I. E., Yu, C., Huszagh, A., Xu, Y., Viner, R., . . . Huang, L. (2017). The proteasome-interacting Ecm29 protein disassembles the 26S proteasome in response to oxidative stress. *J Biol Chem, 292*(39), 16310-16320.

Webb, B., Viswanath, S., Bonomi, M., Pellarin, R., Greenberg, C. H., Saltzberg, D., & Sali, A. (2018). Integrative structure modeling with the Integrative Modeling Platform. *Prot Sci, 27*, 245-258.
